# Revisiting the Use of Structural Similarity Index in Hi-C

**DOI:** 10.1101/2021.09.23.459925

**Authors:** Hanjun Lee, Bruce Blumberg, Michael S. Lawrence, Toshi Shioda

## Abstract

Identification of dynamic changes in chromatin conformation is a fundamental task in genetics. In 2020, Galan et al.^1^ presented CHESS (Comparison of Hi-C Experiments using Structural Similarity), a novel computational algorithm designed for systematic identification of structural differences in chromatin-contact maps. Using CHESS, the same group recently reported that chromatin organization is largely maintained across tissues during dorsoventral patterning of fruit fly embryos despite tissue-specific chromatin states and gene expression^2^. However, here we show that the primary outputs of CHESS–namely, the structural similarity index (SSIM) profiles–are nearly identical regardless of the input matrices, even when query and reference reads were shuffled to destroy any significant differences. This issue stems from the dominance of the regional counting noise arising from stochastic sampling in chromatin-contact maps, reflecting a fundamentally incorrect assumption of the CHESS algorithm. Therefore, biological interpretation of SSIM profiles generated by CHESS requires considerable caution.

---

To reproduce the major findings of Galan et al.^1^, we performed CHESS analysis on the same Hi-C dataset^3^ (diffuse large B-cell lymphoma and healthy B cells) using identical input parameters (Methods). CHESS measures SSIM in every pair of 7-by-7 matrices^4^ in the given window and takes their mean as the main output. Hence, the reproducibility of the results was confirmed by examining the distribution of *z*-normalized mean SSIM (*z*-SSIM) measured by comparing the chromatin-contact maps (Fig. 1a; Pearson’s *r* = 0.87, *P* = 3.1×10^−52^). To assess the validity of the results with respect to the input, we generated random input files by shuffling the reads from the FASTQ files of both cell types (Extended Data Fig. 1), such that each shuffled file has an identical fraction of reads from each sample (Methods). This “sanity check” should have produced an output entirely different from that produced using the bona fide datasets. Alarmingly, the distribution of *z*-SSIM between the shuffled data produced an output nearly indistinguishable from that generated by the real data (Fig. 1a; Pearson’s *r* = 0.87, *P* = 1.4×10^−54^). This observation raises a serious concern that the *z*-SSIM outputs of CHESS are independent of the inputs provided to the software.

**Fig. 1.**
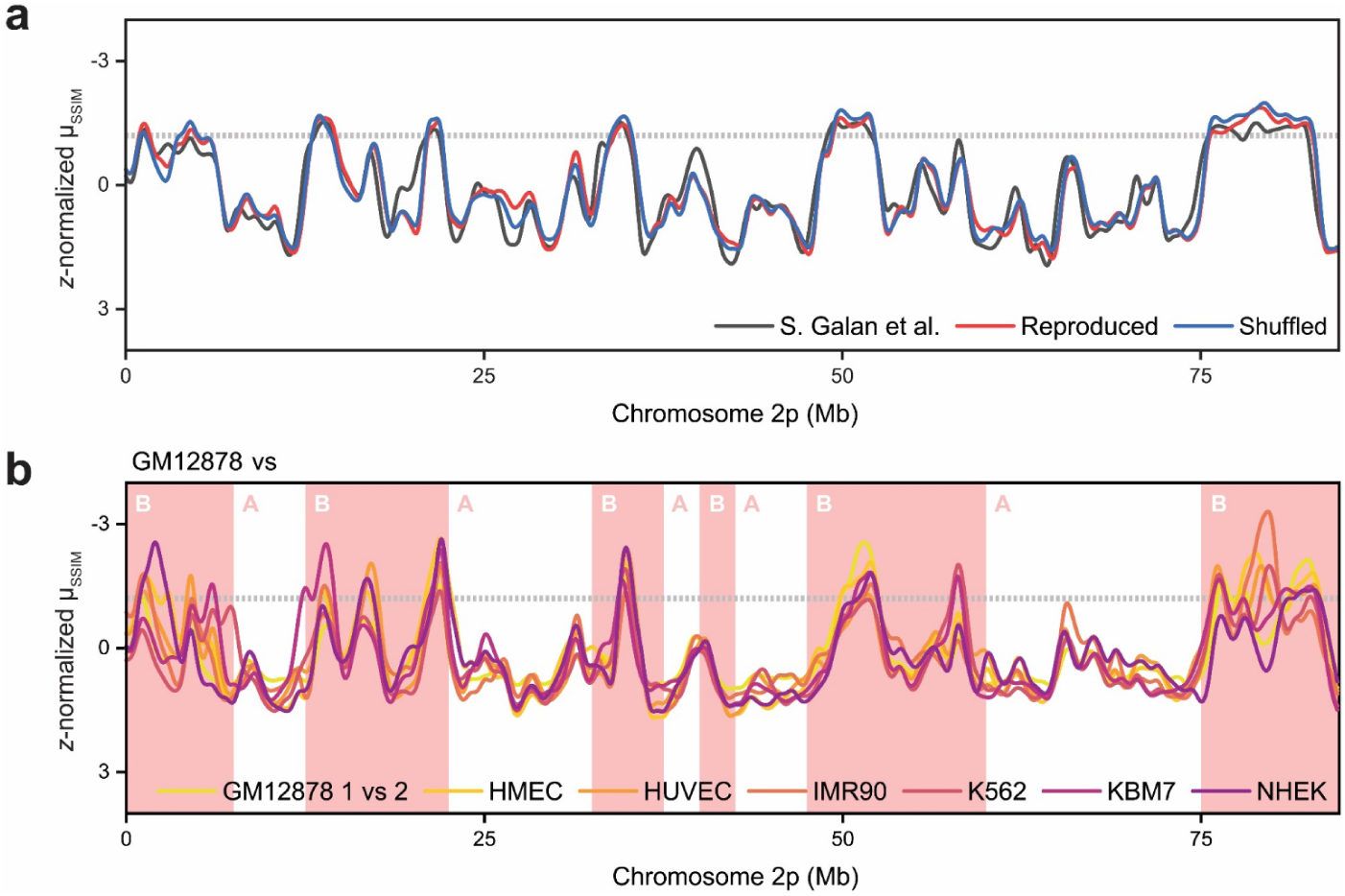
Distributions of mean structural similarity index in Hi-C experiments. **a**, Distributions of *z*-normalized mean structural similarity index (*z*-SSIM) in chromosome 2p of the diffuse large B-cell lymphoma and healthy B cells dataset^3^. The distribution of *z*-SSIM measured by Galan et al.^1^ is shown as a black line, while the reproduced distribution is shown as a red line. To assess the specificity of these distributions to the input matrices, we measured *z*-SSIM for the shuffled files with identical fractions of both the diffuse large B-cell lymphoma and healthy B cells, shown as a blue line. Overall, high correlation was observed between these distributions, suggesting the nonspecific behavior of CHESS. **b**, Distributions of *z*-SSIM in chromosome 2p of seven human cell types^5,6^. Except for the comparison of two biological replicates of GM12878, all cell types were compared against GM12878 B-lymphoblastoid cells. High correlation was observed between all *z*-SSIM distributions. Nuclear compartments measured from GM12878 datasets are indicated in pink and white. The great majority of regions with low *z*-SSIM were located in compartment B. Dotted lines indicate the CHESS threshold for *z*-SSIM significance.

The above concern was corroborated by the *z*-SSIM outputs generated by CHESS on a publicly available Hi-C dataset consisting of seven human cell types^5,6^ (GM12878, HMEC, HUVEC, IMR90, K562, KBM7, and NHEK). Among these data, the chromatin-contact map of the human GM12878 B-lymphoblastoid cells showed the highest density^5^ and hence was used as the control. Again, very strong correlations were observed among the *z*-SSIM distributions of these cell types (Fig. 1b; Pearson’s *r* = 0.80; 95% confidence interval, 0.76–0.84). Even when comparing the Hi-C matrices from two biological replicates of GM12878, we obtained a nearly identical distribution of *z*-SSIM (Fig. 1b; Pearson’s *r* = 0.79; 95% confidence interval, 0.76– 0.83). To completely rule batch effects out, we also performed data shuffling on FASTQ reads of a single Hi-C experiment (Methods) and observed a very strong positive correlation between its *z*-SSIM output and the *z*-SSIM distribution of Galan et al.^1^ (Extended Data Fig. 2; Pearson’s *r* = 0.71, *P* = 7.0×10^−27^). When identical data files are provided as both query and reference, all mean SSIM values became one as previously reported^1^. Overall, our sanity check procedures demonstrated an objectionable behavior of CHESS–that its SSIM outputs are largely independent of the input matrices.

While investigating the potential source of error in CHESS, we noticed that the great majority of peaks with the smallest mean SSIM were located in compartment B (Fig. 1b; *P* = 2.5×10^−5^, Fisher’s exact test), which generally shows lower read coverage depth and more prominent regional noise^7-9^. To assess the impact of the regional noise generated by low read coverage, we focused on the comparison of K562 chronic myelogenous leukemia cells to GM12878 B-lymphoblastoid cells (Fig. 2a). CHESS analysis was performed using the published thresholds for *z*-SSIM and the Fano factor (we note that Galan et al.^1^ used the term signal-to-noise ratio against its conventional definition^10^ to describe the inverse of the Fano factor). Indeed, a significant positive correlation was observed between the logarithm of the read coverage and the distribution of *z*-SSIM (Fig. 2a; Pearson’s *r* = 0.44, *P* = 3.5×10^−5^), indicating that the read coverage is an unexpected determining factor of the intrinsic SSIM profile generated by CHESS.

**Fig. 2.**
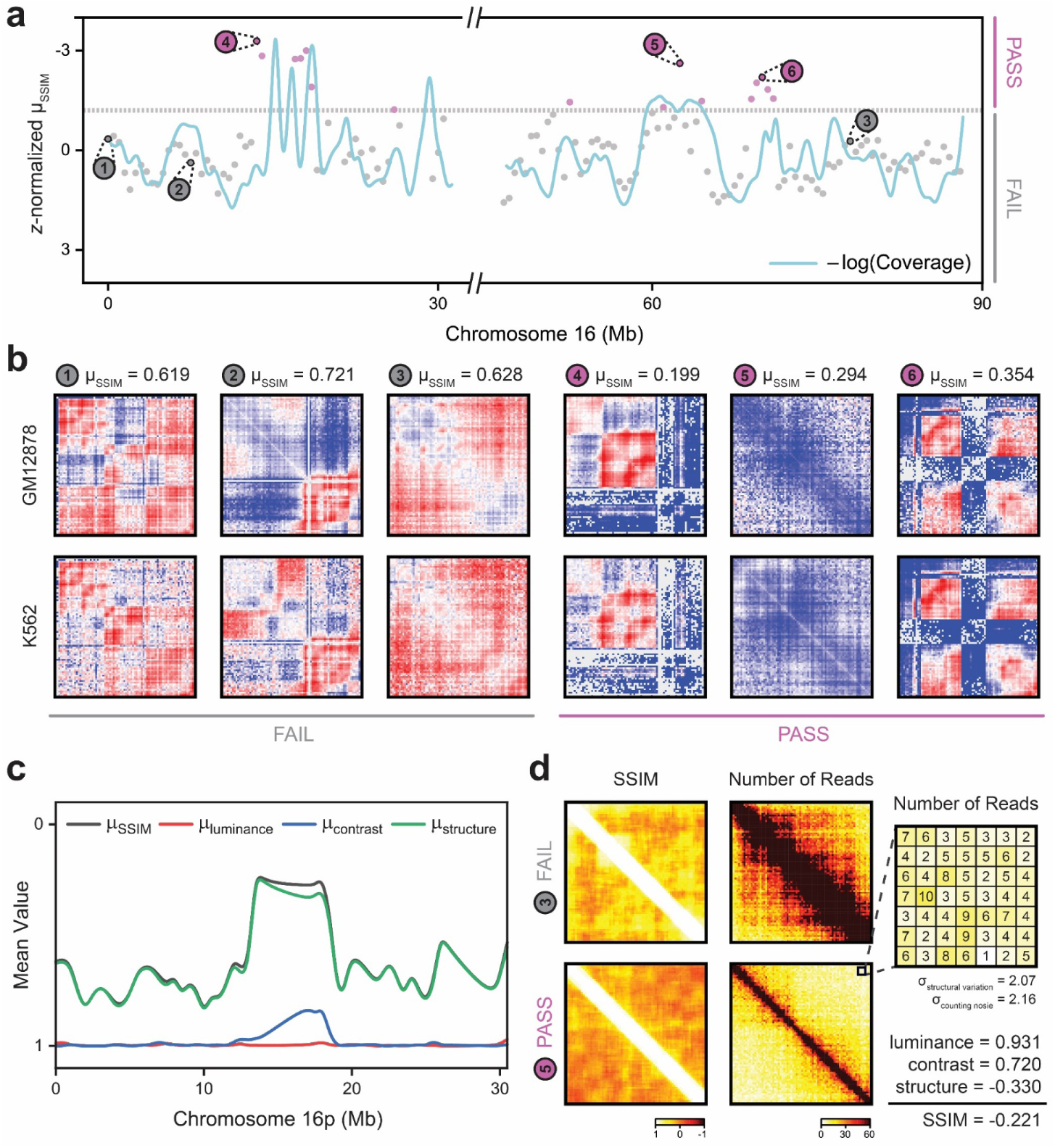
Intrinsic limitations of applying structural similarity index in Hi-C experiments. **a**, Distributions of *z*-normalized mean structural similarity index (*z*-SSIM) in chromosome 16 of K562 chronic myelogenous leukemia cells compared against GM12878 B-lymphoblastoid cells. Dotted line indicates the CHESS threshold for *z*-SSIM significance. Regions that passed both of the CHESS thresholds (i.e., z-SSIM and Fano factor) are indicated in magenta, while those that failed to pass at least one of the thresholds are indicated in gray. For each group, three representative regions were selected for further analyses (panels 1–6). The negative logarithm of read coverage is shown as a blue line, and it strongly resembles the *z*-SSIM distribution. **b**, The observed/expected matrices for representative regions that passed (magenta) or failed to pass (gray) the CHESS thresholds. Matrices for both the GM12878 and K562 datasets are presented. Mean structural similarity index for each comparison is shown in the top lane. **c**, Component analysis of the mean structural similarity index in chromosome 16p of K562 against GM12878. The mean values for luminance (red), contrast (blue), and structure (green) are shown. The structure component had the strongest positive correlation with the distribution of mean structural similarity index (gray). **d**, Stochastic counting noise in chromatin contact maps derived from Hi-C experiments. The distribution of raw structural similarity indices and the number of reads per pixel for K562 are presented for panels 3 and 5. Low similarity indices mainly arose from regions with low read coverage depth. For the 7-by-7 window in panel 5 with the most negative raw structural similarity index, we show the reads-per-pixel matrix. The individual components of the similarity index are shown at bottom right. Overall, our data indicate that the intrinsic counting noise arising from stochastic sampling overwhelms the actual structural variations in determining the value of the structural similarity index.

To assess whether regions with lower mean SSIM exhibited more striking differences, we extracted the observed/expected matrices from regions that passed or did not pass the CHESS thresholds for *z*-SSIM and the Fano factor (Fig. 2b). For regions that passed all of the thresholds set by Galan et al.^1^, those with the local minimum of *z*-SSIM were selected. All three regions that passed the thresholds did not show evidence of differential chromatin interaction, but rather contained significant amount of noise arising from alignment ambiguities and low read coverage (Fig. 2b, panels 4–6). In contrast, there were other regions that failed to pass the CHESS thresholds but exhibited more striking differences in the chromatin-contact map, which included the HBA1 locus previously identified by other computational algorithms^11^ (Fig. 2b, panels 1–3). These results also suggest that the threshold for the Fano factor failed to rescue CHESS from its fundamental error of applying SSIM to Hi-C matrices.

SSIM is defined as the product of components representing similarities in the luminance, contrast, and structure of the image^1,12^. The covariance term included in the structure component is particularly susceptible to variation from regional noise and may explain the abnormal behavior of the similarity index in analyzing chromatin-contact maps. To test this hypothesis, we measured mean values for each component of the similarity index throughout chromosome 16p (Methods). Indeed, the distribution of mean structure strongly resembled the distribution of mean SSIM (Fig. 2c; Pearson’s *r* = 1.0, *P* = 2.4×10^−37^).

To assess whether the read coverage in these Hi-C matrices is low enough to strongly affect the structure component of the similarity index, we examined the distribution of raw SSIM as well as the number of reads in regions that passed or did not pass the CHESS thresholds. Indeed, a striking difference in the number of reads was observed between regions that passed and regions that failed to pass the thresholds. Raw SSIM was the smallest in regions that had the least number of reads per pixel (Fig. 2d). For a 7-by-7 window that had the lowest raw SSIM, the average number of reads per pixel in the K562 data was mere a 4.7, with a window-wide standard deviation of 2.07 (Fig. 2d). Modeling DNA sequencing as a Bernoulli trial of capturing chromatin contacts from a pool of cell nuclei^13^, the standard deviation that arises merely from counting noise becomes 2.16, which is even greater than the window-wide standard deviation in the given matrix. This indicates that the intrinsic noise generated by the low read coverage in these Hi-C matrices overwhelms actual structural variations in determining the value of SSIM.

In summary, we show that there is an intrinsic limitation in the use of SSIM generated by CHESS and that it arises from fundamental limitations in read coverage. As image analysis has become an increasingly popular framework to identify changes in chromatin conformations, we urge researchers to be mindful of the prevalence of regional noise in these data. We also strongly recommend the use of sanity check procedures, such as data shuffling, in bioinformatics analyses, considering the ever-expanding impact of computational approaches.

## Methods

### The CHESS pipeline

The CHESS (v.0.3.7) software was installed directly from the Python Package Index (PyPI) with the following dependencies: Cython (v.0.29.23), FAN-C (v.0.9.20), Future (v.0.18.2), IntervalTree (v.2.1.0), Kneed (v.0.7.0), NumPy (v.1.21.1), Pandas (v.1.3.0), Pathos (v.0.2.8), PyBEDTools (v.0.8.2), Scikit-Image (v.0.18.2), SciPy (v.1.7.0), and Tqdm (v.4.61.2). All chromatin-contact maps were binned at 25-kb resolution and were normalized using Knight-Ruiz (KR) matrix balancing^14^ before analysis. The distribution of *z*-normalized mean structural similarity index (*z*-SSIM) was calculated using the sim functionality of the software. Following input parameters were used for the partitioning of the human hg19 genome and the subsequent calculation of *z*-SSIM: 2-Mb windows span and 500-kb step size. Regions with *z*-SSIM less than or equal to −1.2 and the Fano factor (i.e., window-wide variance divided by window-wide mean) less than or equal to 0.6^-1^ were considered as passing the CHESS thresholds^1^.

### Generation of chromatin-contact maps

HIC files for both the diffuse large B-cell lymphoma and healthy B cells datasets^3^ were directly downloaded from the GitHub repository (https://github.com/vaquerizaslab/chess/tree/master/examples/dlbcl). For the reproduction of the data, we downloaded the raw FASTQ reads directly from the ArrayExpress Archive (accession code, E-MTAB-5875). The first halves of FASTQ files from both samples were merged to generate the first shuffled FASTQ file. Similarly, the second halves of FASTQ files were merged to generate the second shuffled FASTQ file. Each shuffled FASTQ file contained the following number of reads from each file: ERR2704804, 126,115,662 reads; ERR2704805, 125,440,698 reads; ERR2704806, 77,570,082 reads; and ERR2704807, 75,313,370 reads. Overall, both shuffled FASTQ files had an identical fraction of reads from both the diffuse large B-cell lymphoma and healthy B cells (Extended Data Fig. 1). FASTQ files were converted to chromatin-contact maps by running HiC-Pro (https://github.com/nservant/HiC-Pro) using default parameters. Chromatin-contact maps without ICE normalization were utilized for future analyses. HIC files for seven human cell types^5,6^ were directly downloaded from the Gene Expression Omnibus (accession code, GSE63525). Chromatin-contact maps with a MAPQ score greater than or equal to 30 were utilized for analyses. Data shuffling on a single Hi-C experiment was done by splitting the raw FASTQ files downloaded from the Gene Expression Omnibus (accession code, GSM2360314). Each shuffled FASTQ file contained the following number of reads from the original data file: GSM2360314, 80,324,682 reads. For all data shuffling experiments, there was no overlap between the query and the reference.

### Pre-processing of chromatin-contact maps

Chromatin-contact maps binned at 25-kb resolution were extracted and converted to COOL matrices using HiCExplorer (https://github.com/deeptools/HiCExplorer). Read coverage data was retrieved by taking the sum of each column of chromatin-contact maps before normalization. Knight-Ruiz (KR) matrix balancing^14^ was applied to all COOL matrices using the hicConvertFormat functionality of HiCExplorer before running CHESS analyses to match the normalization method of Galan et al.^1^

### Individual analysis of the structural similarity index

Component analysis of the structural similarity index was performed by directly modulating the structural_similarity function of NumPy (v.1.21.1). The following mathematical definitions^12^ for each component of the structural similarity index were used:

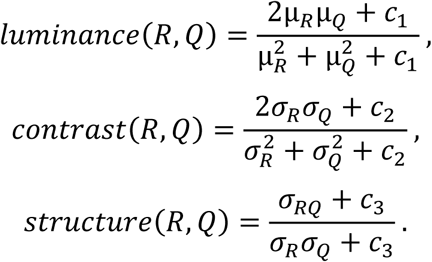

The source codes for these python functions are publicly available at https://github.com/hanjunlee21/StructuralSimilarity/tree/main/ComponentAnalysis. Raw values of the structural similarity index were measured using a custom MATLAB (version R2021a) algorithm publicly available at https://github.com/hanjunlee21/StructuralSimilarity/tree/main/MATLAB. The observed/expected matrices were prepared from the COOL matrices using HiCExplorer and was subsequently converted to GenomicInteractions matrices. For a 7-by-7 window, the raw value of the structural similarity index was calculated using the following definition of the structural similarity index^12^:

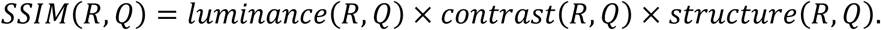

### Estimation of the counting noise

Modeling DNA sequencing as a Bernoulli trial of capturing chromatin contacts from a pool of cell nuclei, the coefficient of variation in the raw count of reads can be approximated as the inverse of the square root of the average number of reads^13^. As a consequence, the standard deviation arising from counting noise can be estimated as the square root of the average number of reads per pixel in the given 7-by-7 window.

### Measurement of the eigenvector

Eigenvectors, required for the determination of the nuclear compartment^7^, were calculated using the eigenvector functionality of Juicer Tools (https://github.com/aidenlab/juicer). Eigenvectors were calculated at a 2.5-Mb resolution with KR matrix balancing^14^. Eigenvectors were compared against the publicly available DNase I hypersensitivity assay dataset for GM12878 B-lymphoblastoid cells (ENCODE accession code, ENCSR000EMT) to fully determine the sign of the element of the eigenvector for each compartment.

### Statistics

All statistical analyses were performed on OriginPro (version 2021b). For the correlation analysis of *z*-SSIM distributions, Pearson’s correlation coefficient was assessed. Group of Pearson’s correlation coefficients from multiple *z*-SSIM distributions were presented as their mean and their 95% confidence intervals in the manuscript. Fisher’s exact test was used for a two-sample proportion test.

## Reporting Summary

Further information on research design is available in the Nature Research Reporting Summary linked to this article.

## Data availability

HIC files for both the diffuse large B-cell lymphoma and healthy B cells datasets^3^ are available for download at https://github.com/vaquerizaslab/chess/tree/master/examples/dlbcl, while the raw FASTQ files can be accessed from the ArrayExpress Archive under the accession code E-MTAB-5875. COOL files for the reproduced and shuffled data can be accessed from https://github.com/hanjunlee21/StructuralSimilarity/tree/main/COOL. HIC files for seven human cell types^5,6^ are available for download at the Gene Expression Omnibus under the accession code GSE63525. DNase I hypersensitivity assay dataset for GM12878 is available for download at https://www.encodeproject.org/experiments/ENCSR000EMT/.

## Code availability

All codes required for the reproduction of our finding is available on GitHub (https://github.com/hanjunlee21/StructuralSimilarity). The CHESS source code^1^ is publicly available at https://github.com/vaquerizaslab/CHESS and is indexed in the Python Package Index (PyPI) as chess-hic.

## Acknowledgements

We thank the support of the National Institute of Health (grant no. R01ES023316 and R01ES031139 to B.B. and T.S.). H.L. thanks the support from the Kwanjeong Educational Foundation Overseas Scholarship Program.

## Contributions

B.B. and T.S. conceptualized the study. H.L. carried out the investigation. H.L. prepared and wrote the original draft of the manuscript. H.L., B.B., M.S.L., and T.S. reviewed and edited the draft. M.S.L. and T.S. supervised the study.

## Ethic Declarations

### Competing Interests

The authors declare no competing interests.

## Notes

https://github.com/hanjunlee21/StructuralSimilarity

### Competing Interest Statement

The authors have declared no competing interest.

