## Supplementary Figures for "Revisiting the Use of Structural Similarity Index in Hi-C"

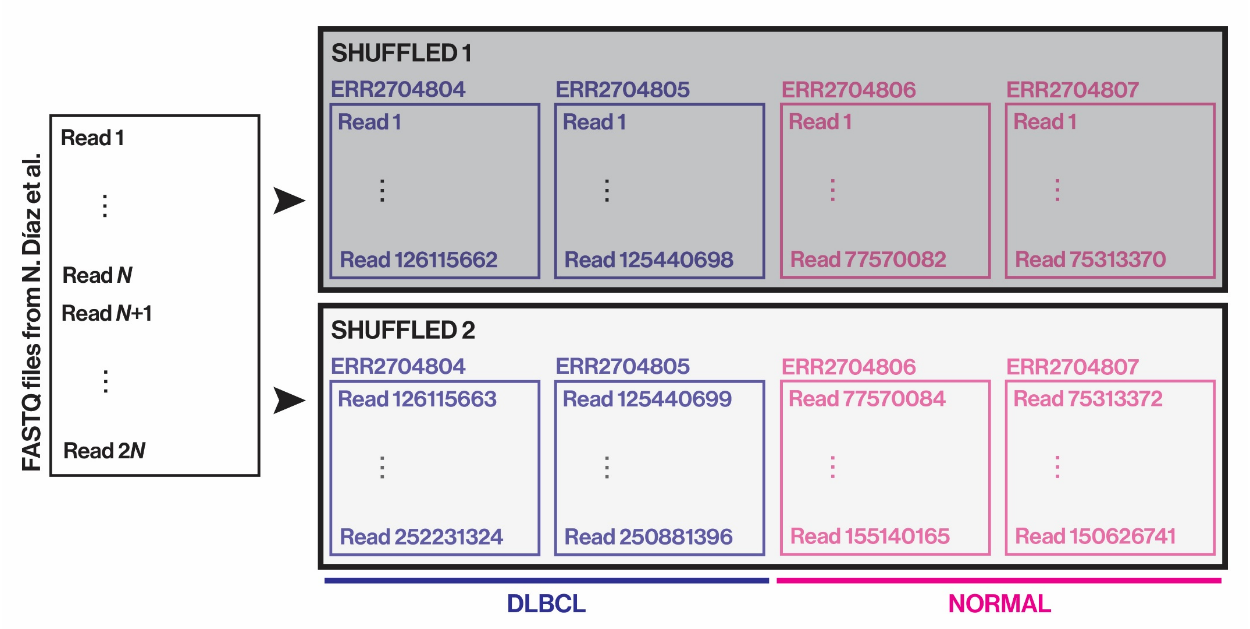


**Extended Data Fig. 1 | Schematic of data shuffling.** To destroy any significant differences in chromatin contacts between the query and reference input files of CHESS, reads from the FASTQ files of Díaz et al.[3] were shuffled to create hybrid FASTQ files containing identical fraction of reads from DLBCL and NORMAL libraries. DLBCL, diffuse large B-cell lymphoma.

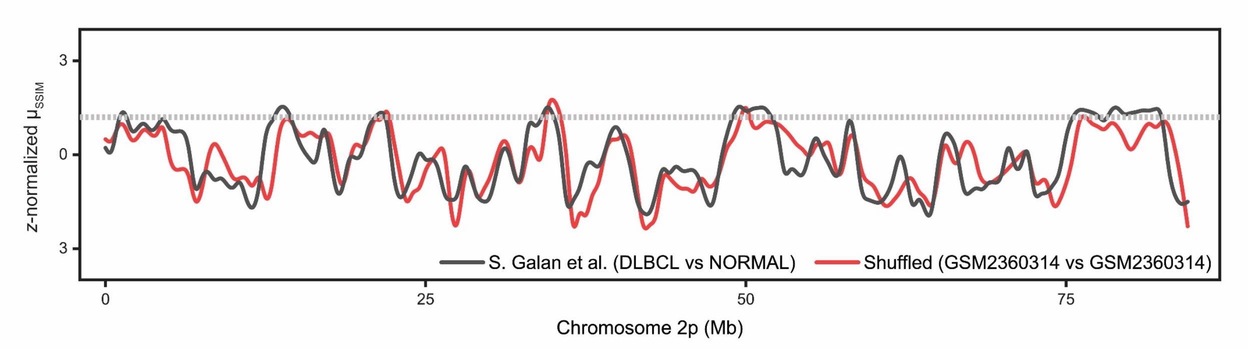


**Extended Data Fig. 2 | Data shuffling on a single Hi-C experiment.** To completely rule batch effects out, reads from a single Hi-C experiment (GEO accession code, GSM2360314) were randomly extracted to generate two shuffled FASTQ files. Very strong positive correlation was observed between the z-SSIM output using shuffled data (red) and the z-SSIM distribution of Galan et al.[1] (black), despite originating from different Hi-C datasets. Dotted line indicates the CHESS threshold for *z*-SSIM significance.
